# Characterizing circadian connectome of *O. tenuiflorum* using an integrated network theoretic framework

**DOI:** 10.1101/2022.03.02.482599

**Authors:** Vikram Singh, Vikram Singh

## Abstract

*Ocimum tenuiflorum* (Tulsi) is among the most valuable medicinal plants as almost every part of this herb and the essential oil it produces possess pharmaceutical properties that have been utilized since ancient times to cure a large number of diseases. Like in other plants, circadian clock in Tulsi regulate important physiological processes like growth, development, defence *etc*. by anticipating environmental cues. In the present work, identification and characterization of core circadian clock and clock associated proteins in Tulsi genome is reported. We mined 24 core clock (CC) proteins in 56 template plant genomes to build their hidden Markov models (HMMs). These HMMs were used to identify 24 core clock proteins in *O. tenuiflorum* which were further subjected to functional annotation. A hybrid network theoretic methodology comprising of random walk with restart (RWR) and graphlet degree vectors (GDV) was implemented to explore the local topology of the interologous, Tulsi protein interaction network (TulsiPIN) and mine CC associated raw candidate proteins. Statistical and biological significance of the raw candidates was determined using permutation and enrichment tests. A total of 70 putative CC associated proteins were identified which were further subjected to functional annotation.

## 1 Introduction

The earth’s rotation around its own axis and revolution around the sun generates repetitive changes in the environmental conditions, like, dark-light cycle, rhythmicity of light intensity, temperature, seasons etc. Guided by these environmental rhythms, in living organisms exist biological rhythms called circadian rhythms which are endogenously generated, self sustained and temperature compensated [2, 31, 57]. These rhythms directly influence the physiological processes of all the living beings including plants that are helpful in modulating and temporal organisation of biological traits [15]. The involvement of Circadian clock in regulating vital plant processes, like, nutrient biogenesis, root growth, photosynthesis, hormonal signalling, sugar metabolism, flowering time, and plant immunity has been documented in a large number of studies [10, 66, 11, 65, 28]. These biological rhythms are entrained to their external environment by an internal time keeper called circadian clock that keeps track of environmental fluctuations and allow temporal separation of internal biological events accordingly [18]. Circadian clock is ubiquitous and has a period approximately equal to 24 hours that persists in free running conditions *i*.*e*. in the absence of any external cues [17]. Circadian clock is crucial for optimizing plant’s adaptive responses, in different physiological processes including growth and development, to varying environmental conditions and improve overall fitness of the plant [69, 11]. Circadian oscillators are ubiquitous among all the three domains of life with a very little elemental (gene) conservation across them, however, their architectures and mechanisms are remarkably convergent across domains [32, 14, 13].

All plant circadian clocks anticipate time with the help of input pathways which relay time-encoded information to entrain the central oscillator [19]. Finally the output of the core oscillators is transmitted through output pathways that allow temporal regulation of plant physiological processes [15]. The central oscillator that generates these rhythms is comprised of about 20 genes which are regulated by interlocked transcriptional and transclational feedback loops [41]. At dawn, morning phased transcription factors CIRCADIAN CLOCK ASSOCIATED 1 (CCA1) and LONG ELONGATED HYPOCOTYL (LHY) genes downregulate the expression of afternoon phased genes including PSEUDO-RESPONSE REGULATOR 5 (PRR5), PRR7, PRR9, TIMING OF CAB EXPRESSION 1 (TOC1/PRR1). Other direct targets of CCA1/LHY complex include evening phased genes GIGANTEA (GI) and genes belonging to evening complex (EC), namely, LUX ARRHYTHMO (LUX), EARLY FLOWERING 3 (ELF3), ELF4 [41]. On the other hand morning and midday phase genes including TOC1 along with other PRR family genes and CCA1 HIKING EXPEDITION (CHE) subsequently express during day and provide negative feedback by restricting CCA1 and LHY expression during day time. At afternoon, REVEILLE 8 (RVE8) and RVE4 along with NIGHT LIGHT-INDUCIBLE AND CLOCK-REGULATED GENE 1 (LNK1) and LNK2 transcription factors have been shown to induce expression of several clock genes including of PRR5, TOC1 and ELF4 genes [41, 28]. In the evening time, TOC1 negatively regulates GI, LUX, ELF4 and PRR5 [28]. The negative regulation of GI is maintained by GI that also downregulates the expression of PRR7 and PRR9.

*Ocimum tenuiflorum* (Tulsi), a member of family Lamiaceae, is an aromatic herb that is widely recognized for array of health benefits it offers [9]. We find references of the holy basil in Ayurveda, an ancient medicinal system, for its uses to treat variety of diseases and disorders either independently or as formulation in combination with several other herbs [44]. The plant is reared for its oils and every part of plant produces a number of secondary metabolites which possess significant pharmaceutical properties [42]. *Ocimum* leaves are reported to possess anthelmintic properties *e*.*g*. prevent ring worm infection, expectorant to alleviate cough, diaphoretic [54] and several imperative medicinal properties like, neuroprotective, memory enhancement, anticonvulsant, analgesic, anticancer, antidiabetic, antistress, antispasmodic, antiulcer, antioxidant, antiinflamatory, antimicrobial [37].

Recently genome [1, 48], tanscriptome [49] and proteome [58] wide studies have reported a draft catalogue of genes, transcripts and proteins present in this species along with important mechanistic insights about secondary metabolite biosynthetic pathways. However, an exhaustive systems level study presenting core circadian clock genes and complex interplay among them at molecular level is still lacking in *O. tenuiflorum*. In this study, we have adopted a systems level approach to mine and characterize core circadian clock proteins giving rise to these rhythms in *O. tenuiflorum*. We leveraged RWR and GDV methods to explore the local topology of protein interaction network of Tulsi (TulsiPIN) and identified raw candidate proteins associated with the core oscillator. Furthermore, statistical and biological significance of the candidates was determined and their functional annotations were determined.

## 2 Materials and methods

### 2.1 Data collection

From the literature 24 core circadian clock proteins of *Arabidopsis thaliana* [41, 28] were identified and their sequences were retrieved from the TAIR database [25]. Further 56 template plants for which genome information is available were enlisted and their proteome data was downloaded from UniProt database https://www.uniprot.org/. All the CC proteins were then queried against each template using Blast-P to identify orthologous proteins. Top hits with at least 40% sequence identity, 50% query coverage, and an *E-value* ≤ 10^−5^ were selected for every sequence and aligned using Clustal Omega [53]. HMMER http://hmmer.janelia.org/ was used to construct Hidden Markov models for each CC protein. These HMMs were used to scan *O. tenuiflorum* proteome obtained from TulsiDB http://caps.ncbs.res.in/Ote/ to identify putative CC proteins in Tulsi. Each identified CC protein was then subjected to InerProScan v5.48-83 https://www.ebi.ac.uk/interpro/ to identify their respective domains [56]. We collected protein-protein interaction network of *O. tenuiflorum* called TulsiPIN from an earlier study [58]. TulsiPIN is an exhaustively constructed network consisted of 13, 604 nodes and 327, 374 edges that possess both scale free and small world properties.

### 2.2 Mining CC associated proteins using local topological measures

Two local network topology based algorithms namely random walk with restart (RWR) and graphlet degree vector (GDV) have been leveraged to identify core clock (CC) associated proteins.

#### 2.2.1 Random walk with restart (RWR) based mining

RWR is a ranking algorithm that has been successfully applied to estimate the likelihood of genes/proteins being associated with a given biological process (X) based on their proximity to validated proteins belonging to X in the input network [24]. The algorithm initiates walks from one or more seed nodes (CC proteins) to some random node(s) in the network and returns the probability of each protein being a core clock associated gene. In this study RWR algorithm was applied on TulsiPIN using 9 CC proteins (which could map to TulsiPIN) as seed nodes. Initially a vector *P*_0_, containing *M* elements each corresponding to a node in TulsiPIN, was constructed in which each CC genglobale was initialized with an equal value of 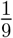 while other *O. tenuiflorum* genes with a value of zero. The vector is then updated using the equation *P*_*i*+1_ = (1 − *r*)*A*^*T*^ *P*_*i*_ + *rP*_0_ for *i*^*th*^ iteration until *P*_*i*_ is stabilized. Here *r* represents restart probability that is set as (0.8), and *A* is TulsiPIN’s adjacency matrix, every column of that sums up to one. The iterations stop when Manhattan distance between *P*_*i*_ + 1 and *P*_*i*_ is less than 10^−6^ *i*.*e*. 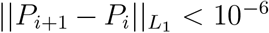. This yielded a probability vector containing the probability values corresponding to each protein of TulsiPIN quantifying their likelihood of being CC associated. Accordingly a threshold value of 10^−5^ was adopted and all the proteins having probabilities higher than the selected threshold called CC associated proteins were considered for further study [7].

#### 2.2.2 Graphlets degree vector (GDV) based mining

Small, non-isomorphic induced sub-graphs of a large network are defined as graphlets[47]. Based on the node position within a graphlet, each subgraph be differentiated into symmetrically equivalent sets of nodes (symmetry groups) called automorphism orbits [46]. A total of 15 automorphism orbits are possible for up to 4 node graphlets and each node of the network can be represented with a vector of length 15, each element of which represents the degree of corresponding orbit, called graphlet degree vector (GDV). Finally a vector of 11 non-redundant orbits [67] containing orbit degrees, representing wiring around that node, was computed for every node in the network. Since current PPI networks have been shown to be largely incomplete and lower degree nodes have higher likelihood of being present in the those incomplete parts so we remove proteins with degree lower than three [35]. Among the 24 CC proteins only 10 could be mapped to TulsiPIN and one of them has degree lower than three so removed from the further study. For each of 9 CC proteins a pairwise similarity (*S*_*GDV*_) was computed with every TulsiPIN protein having degree more than three that is defined as

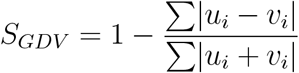

where *u* and *v* represent GDV of CC proteins and TulsiPIN proteins respectively while *i* represents the *i*^*th*^ orbit. All the proteins having similarity above 94% with the protein of interest were considered as core clock associated proteins. Previously a threshold between 90% and 95% have been shown to be optimal [35]. So to pick upper threshold we started with a threshold value of 96% and decreased the threshold value until we obtained at least one TulsiPIN protein similar with each CC protein other than themselves.

### 2.3 Statistical and functional validation of CC associated proteins

As these algorithms exploit the topology of the network to estimate the likelihood, this may output some false positive proteins as CC associated proteins *e*.*g*. hubs will ranked higher than those having degree lower than the average degree of network [7]. To curb these false positive proteins and assess the reliability of each prediction two methods, namely, permutation and enrichment tests were applied.

#### 2.3.1 Permutation or randomisation test

To test the statistical significance of CC associated proteins identified by RWR algorithm, we sampled 1000 random sets of 9 proteins each from *N* TulsiPIN proteins. RWR algorithm was performed on TulsiPIN using each sample as seed. Then a *p-value* or permutation FDR for every CC associated proteins predicted was computed based on the following equation 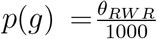, where *θ* is the number of random sets for which probability of *g* is higher than *p*(*g*) obtained using CC proteins as seed. On the other hand to test the statistical significance of CC associated proteins obtained from GDV algorithm we first constructed an ensemble of 1000 random networks with same number of nodes and similar degree sequence [36] and then computed graphlet degree vectors for each protein of every network. Then pairwise *S*_*GDV*_ similarity scores were computed for each of the CC protein with other proteins in TulsiPIN. Similar to RWR, a permutation FDR for every CC associated protein predicted by GDV algorithm was computed using 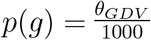, where *θ* is the number of random networks for which the similarity score of protein pair between *g*, a CC associated gene, and some CC gene is higher than the similarity score obtained for same pair in real network. Higher *p-value* of a gene indicates the gene is less likely to be associated with CC. Thus CC associated proteins with *p-value* less than 0.05 were selected, from both methods, and the selected protein were called candidate proteins for convenience.

#### 2.3.2 Enrichment test

Linkage test is based on the premise that interacting proteins are more likely to share common functions [58, 55]. So, in this section, we used PPI information form TulsiPIN and Gene Ontology (GO) and KEGG Orthology (KO) annotation information of interacting proteins to infer the validity of candidate CC associated proteins. It has been shown earlier that similar proteins may perform similar molecular functions, localize in same cellular milieu and may be involved in similar biological processes [58, 55]. Thus candidate proteins having similar functional annotations as that of CC proteins may be novel CC associated proteins. To quantify the relationship between a protein and an annotation terms (GO or KEGG pathways), ontology enrichment score that is the negative log of probability value obtained from hypergeometric test was computed [6]. Given *N* total TulsiPIN proteins among which *M* are annotated by one annotation term (*A*). If we randomly draw a set of proteins (*H*(*g*)) containing any protein *g* and its immediate neighbors then the probability that at least *m* them are annotated with *A* is given by

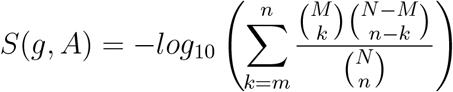

where *n* is the number of proteins present in set *H*(*g*) and *m* is the number of *H*(*g*) proteins annotated by *A*. A vector *ES*(*g*) can be created by computing *S* for all the annotation terms of *g*. These vectors can be used to quantify the similarity between two genes *g* and *g*^t^ as follows

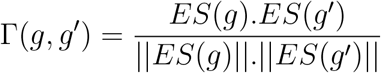

Large value of Γ indicate high functional similarity between the two genes *g* and *g*′. We compute maximum annotation similarity (MAS), that represents the largest Γ value among all possible pairs between a candidate protein *p* and set of CC genes. A threshold value of 0.75 was used to select putative CC associated proteins.

### 2.4 Functional annotation of clock associated proteins

Each protein identified to be associated with core clock of *O. tenuiflorum* was then queried against NCBI non-redundant (nr), UniProt [4] and TAIR [5] databases using Blast-P [3] program to assign putative functions. Default parameters except *E-value*, that is set to be 10^−5^, were used to assign homology. AgriGo [12] and WEGO [68] were used to predict Gene Ontology (GO) terms for each clock associated protein. KEGG ontology (KO) terms were predicted for each clock associated protein using KEGG [22] database. All the 13 programs aggregated in InterProScan v5.48-83 were utilised to predict protein family, functional domains and sites in each clock associated protein [21].

## 3 Results and Discussion

*O. tenuiflorum* plants have been reported to produce secondary metabolites (essential oils) which are mixtures of diverse aromatic compounds. These chemical compounds possess enormous medicinal properties that are being exploited continuously since ancient times [9]. Under varying environmental conditions, plants are subjected to continuous stress either biotic or abiotic that have been shown to hamper plant growth, development and have significant impact on the oil production [50, 43]. Plants’ growth and development is under circadian control. A circadian clock tuned precisely plays substantial role in increasing plants’ fitness and survival by temporally coordinating various vital processes, like, metabolism, physiology *etc*. in accordance with the external conditions [28]. Thus, we attempt to identify core circadian clock and associated proteins in *O. tenuiflorum* through an exhaustive computational framework presented in Figure 1.

**Figure 1:**
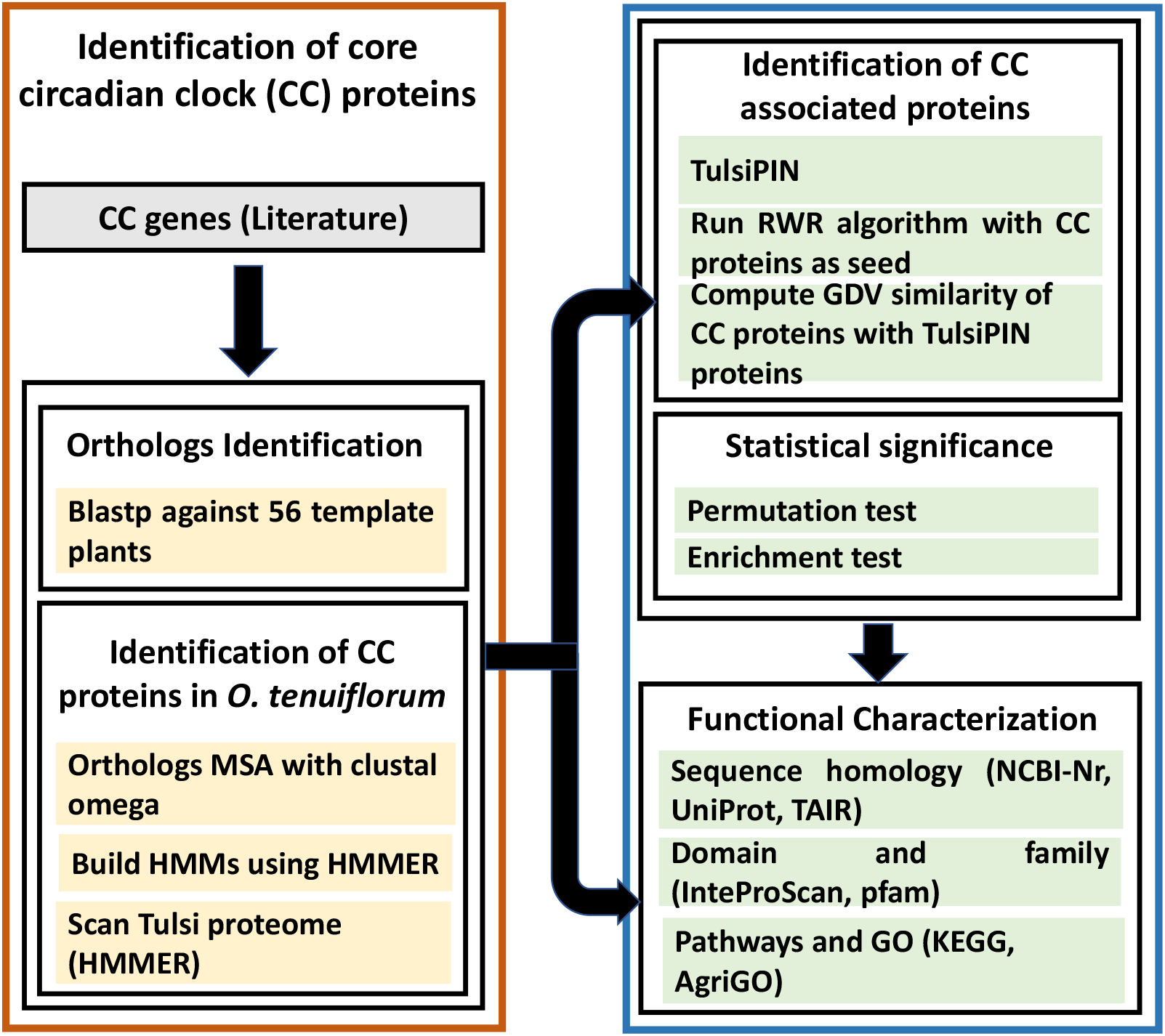
Overall methodology for genome wide identification, characterization of core circadian clock (CC) and CC associated proteins in *O. tenuiflorum*

### 3.1 Mining of core clock proteins and their functional annotation

From literature we identified 24 core circadian clock proteins and attempt to identify them in *Ocimum tenuiflorum*. Proteomes of 56 template plants were scanned using Blast-P for putative ortholgs of selected clock proteins that resulted in 2, 543 hits at an *E-value <* 10^−5^. Further top hits with sequence identity < 40% and query coverage < 50% were filtered out resulting in 963 hits. Separate HMM profiles for each of the core clock protein were built by extracting orthologous sequences of that protein from respective template plants which satisfy the above criteria. These HMMs were used to scan *O. tenuiflorum* proteome based on top hits, all the 24 putative core clock proteins were identified (Supplementary Table S1). Since, transcription factors CCA1 and LHY are partially redundant, transcription regulator LWD1 with LWD2, LUX with NOX, RVE6 with RVE8 and they express during different times of the day or form complexes [41] so we selected the second best hit for LWD2, RVE4, PRR5, RVE6, NOX, PRR3 and LHY proteins (Fig 2).

**Figure 2:**
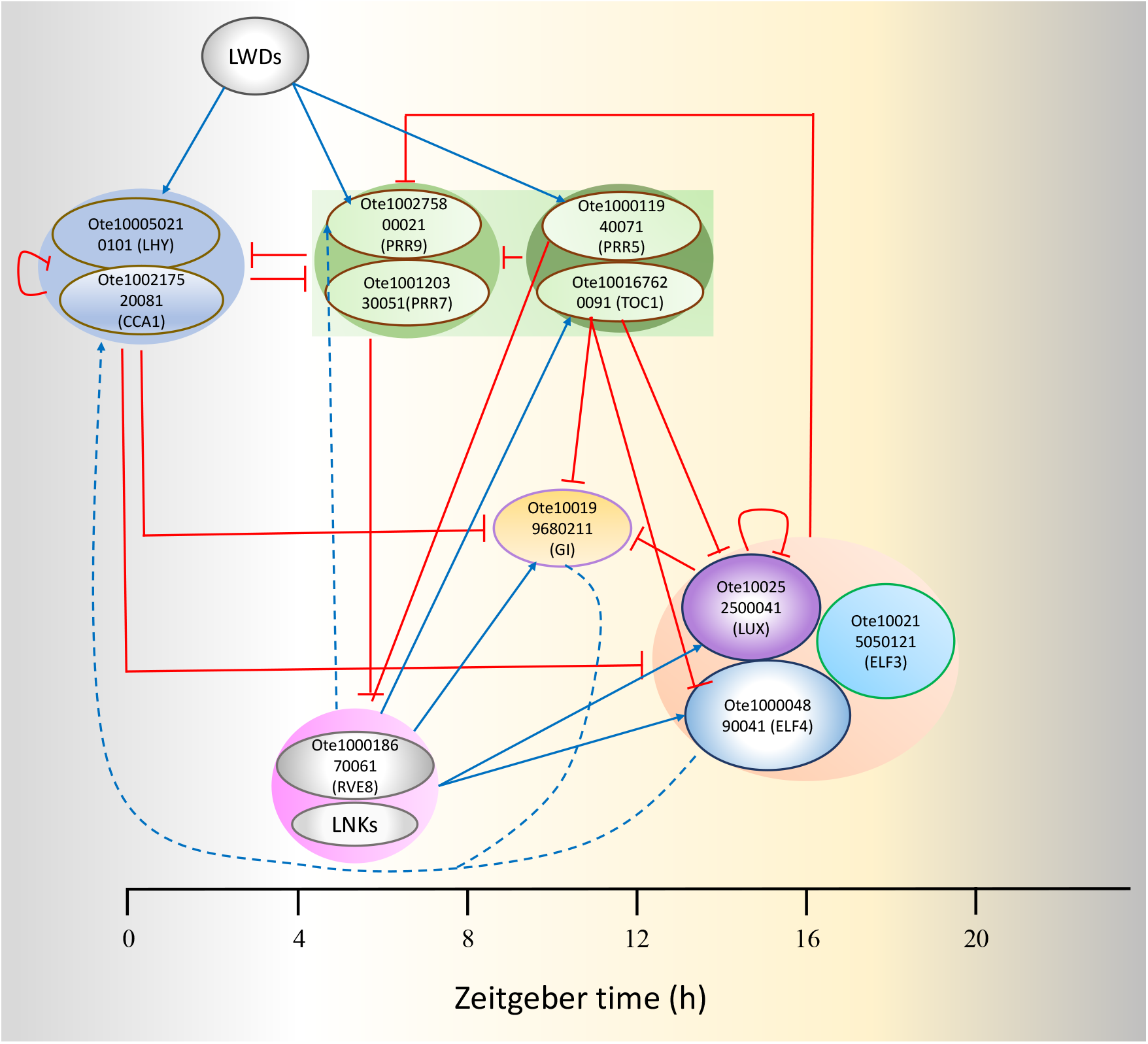
The core clock module of *O. tenuiflorum*

Functions of all the 24 CC proteins were predicted by querying them against NCBI non-redundant, UniProt and TAIR databases using using Blast-P. The results also confirmed the identities of predicted clock proteins and show that most of the proteins possess their respective functions (Table S2). Furthermore, protein family and domain databases integrated by InterProScan also confirmed our results (Table S2). Functional annotation of CC proteins is followed by their gene ontology (GO) annotations (Table S2). A total of 86 GO terms were successfully identified for these CR proteins; with 67 terms classified into 13 categories of biological processes, 2 terms were classified into 2 categories of cellular components and 17 GO terms associated with 2 categories of molecular functions. ‘Regulation of biological process’, ‘Cellular process’, ‘Response to stimulus’ are highly enriched among biological processes followed by ‘signaling’, ‘rhythmic process’. Among molecular functions, ‘binding’ and ‘transcription regulator activity’ were found to be the most enriched categories. Successful assignment of “rhythmic process” associated gene ontology (GO) terms to the identified CC proteins further confirms the role of these proteins as core circadian clock of *O. tenuiflorum*. Furthermore, for finding the CC proteins regulated pathways, KEGG database was used that predicted KO ids of 16 proteins enriching 8 pathways, of these ‘Circadian rhythm in plants’ is found to be most enriched (Table S2). Other pathways include ‘Ubiquitin system’ followed by ‘Plant hormone signal transduction’ and ‘Transcription factors’.

### 3.2 Prediction of circadian clock associated proteins

It is known that different centrality measures address different topological aspects of a network and therefore capture the snapshot of a network in terms of selected architectural metric [29]. For example, connectivity-based measures, assigns higher weights to high degree nodes, on the other hand, middle nodes connecting communities will be ranked higher by measures employing betweenness centrality. So, in this work, two local network topology based algorithms namely random walk with restart (RWR) and graphlet degree vector (GDV) have been leveraged to identify core clock (CC) associated proteins. GDV based approach does not consider only the current node rather it take into consideration the wiring pattern around that node *i*.*e*. it incorporates the information about local neighbourhood. In order to find the circadian clock associated proteins all the identified proteins were mapped to genome wide interologous protein interaction network of Tulsi (TulsiPIN). A total of 10 proteins were successfully mapped to TulsiPIN proteins which were found to interact with 248 other proteins. Since one protein has degree lower than 3 so removed from the study. Remaining nine CC proteins were used as seed nodes to the RWR algorithm applied on TulsiPIN. This produced a probability value corresponding to each protein representing the likelihood of that protein being a novel circadian clock associated protein. All the proteins having *p-value* larger than 10^−5^ were considered for further study resulting in 691 nodes called RWR proteins (Table S3). Because all RWR proteins were not circadian clock related some of them have been selected due to network architecture alone. To filter them out we employed permutation test and computed a *p-value* called the permutation FDR for every RWR protein. All the nodes having probabilities less than 0.05 were selected resulting in 255 putative circadian clock associated proteins called candidate RWR proteins and are presented in Table S3. Similarly we computed pairwise GDV similarity between CC proteins and other TulsiPIN proteins. This resulted in 146 proteins, having *S*_*GDV*_ > 94, predicted to be CC associated by GDV algorithm. Similarly pairwise similarities were obtained from ensemble of 1000 random networks preserving the degree sequence of TulsiPIN. A total of 146 proteins were found to be statistically significant having *p-value <* 0.05 and were called as candidate GDV proteins (Table S3). Furthermore, to add another layer of confidence and select only those proteins which are having similar functions to CC proteins we leveraged Gene Ontology and pathways information to compute the functional similarity (Γ) between each CC protein and every candidate protein. Among 255 RWR candidate proteins a total of 47 were found to have Γ > 0.75 in both GO and pathways information based methods. On the other hand 25 out of 146 proteins were found to have Γ score 0.75 or above in both the methods (Table S3). We combined the proteins obtained from the two methods (RWR and GDV) and found a total of 70 candidate proteins among which 2 common to both the methods. We constructed circadian connectome of Tulsi by mapping the CC and CC associated proteins to TulsiPIN and found that these 94 proteins regulating 2, 412 other proteins and participate in a total of 8, 017 interactions (Fig. 3).

**Figure 3:**
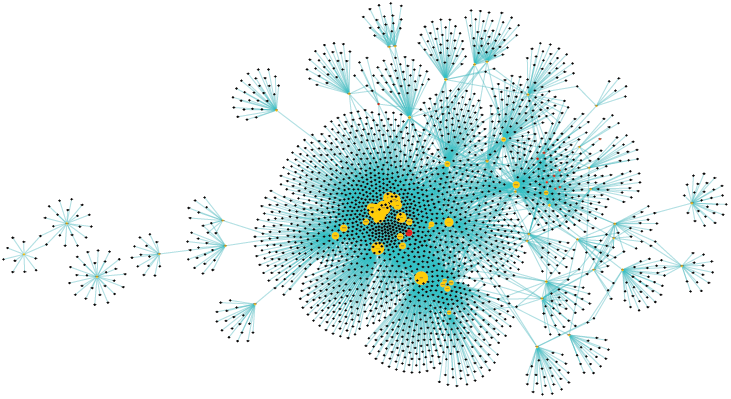
Sub network of proteins obtained by combining core circadian clock (CC) and associated proteins. Yellow colored nodes represent CC associated protenis, red colored nodes are CC proteins

### 3.3 Functional characterization of circadian clock associated proteins

Similar to CC proteins all the 70 CC associated proteins were queried against NCBI non-redundant, UniProt and TAIR databases using using Blast-P, their functions were predicted. The functional annotation of CC associated proteins revealed that most of the proteins are involved in various aspects of growth, development and defence of *O. tenuiflorum* (Table S4). We broadly classified them into seven categories, namely, Gene expression (20), Metabolism, Cell signaling (16), Transport (15), Post translational modification (5), Stress response (2) and Protein folding (1). It has been shown that approximately one-third of genes in *Arabidopsis thaliana* and ≈89% of its transcripts are regulated by varying environmental conditions like temperature, light *etc* [33, 34]. These studies revealed a tight circadian regulation of *A. thaliana* genes at transcriptional level [33] which is consistent with our results that 20 of CC associated proteins are involved in gene expression and regulation (Fig 4). The basic model of circadian regulation is comprised of a core oscillator that is a complex web of multiple coupled transcriptional and translational feedback loops (TTLF). The core oscillator perceive external cues through input pathways and temporally organise various physiological processes of the plant [60].

**Figure 4:**
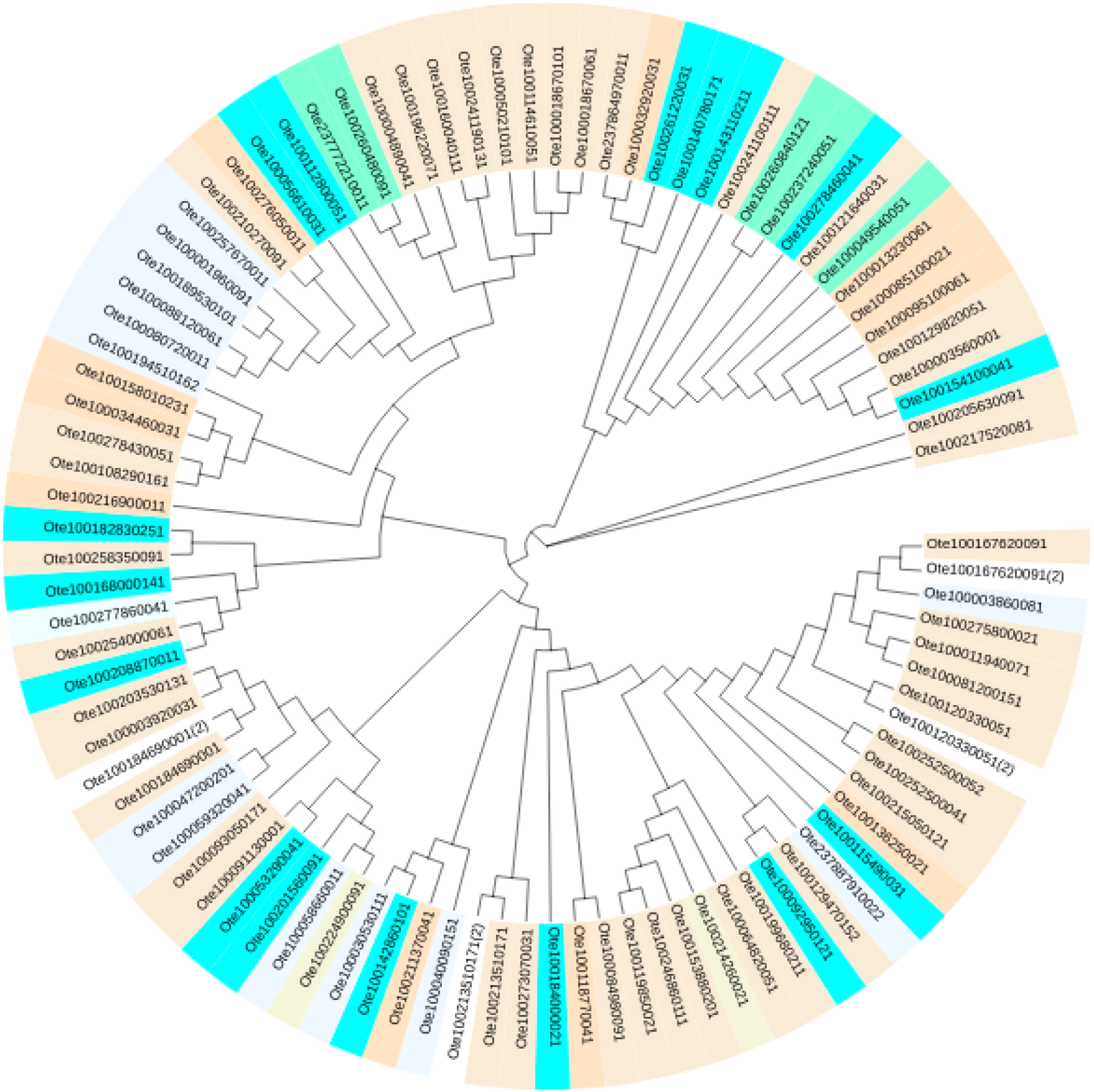
Cladogram of circadian clock circuitry including both CC and associated proteins. Leafs are colored according to their broad function. Cell signaling (aliceblue), Gene expression (antiquewhite), Metabolism (aqua), Post translational modification (aquamarine), Protein folding (azure), stress response (beige), Transport (bisque)

#### 3.3.1 The input module of core clock

Proteins Ote100059320041, Ote100047200201 and Ote10021351017 have been predicted to be light perceiving photoreceptors namely Phytochrome A, Phytochrome B, Adagio protein 1 (ADO1) or ZEITLUPE (ZTL) protein. It is known that Phytochromes primarily perceive red and far red light while cryptochromes and ZEITLUPE (ZTL) family members are blue light photoreceptors. Among the five phytochromes (PHYA-E) in *A. thaliana*, when subjected to continuous red light, PHYA and PHYB have been shown to be primary contributors to photoperception and entrainment of circadian rhythms [59]. Further these signals are transduced to the core oscillator through an intermediate protein CONSTITUTIVE PHOTOMORPHOGENIC 1 (COP1) [52]. Proteins Ote237772210011 and Ote100260480091 have been characterized as an E3 ubiquitin ligase COP1. It triggers light gated proteasomal degradation of core clock components ELF3 and GI hence acts as a direct link between the clock and light perceiving photoreceptors [70, 52]. The expression of ELONGATED HYPOCOTYL5 (HY5), FAR RED ELONGATED HYPOCOTYL3 (FHY3) and FAR RED IMPAIRED RESPONSE1 (FAR1) is under tight regulation of PHYA signaling. During the day they binds directly to ELF4 promoter and induce its expression [26]. The ADO1 and other two members this family possess an F-Box domain, a PAS-like LOV domain and six kelch repeats. F-Box domain links target protein for proteasomal degradation by an E3 Ubiquition ligase complex called skp-cullin-F-Box that is comprised of by interactions between F-Box protein with SKP1, CULLIN, and RBX1 [16]. It has been shown that ADO1 receive input signal by interacting with carboxy terminus of blue light photoreceptors phyB and cry1 [20] and mediates blue light dependent degradation of TOC1 and PRR5 through GI mediated stabilization [23]. Ote100194510162 has been characterized as a serine/threonine phosphatase 5 protein. It has been shown to dephosphorylate a specific serine residue on pfr phytochromes. This dephosphorylation enhances phytochrome stability as well as increase its ability to bind downstream signal transducers and thus enhances phytochrome mediated photoresponses [51]. Proteins Ote237887910022 and Ote100030530111 have been characterized as serine/threonine phosphatase 7 that has been show to implicate in cryptochromes mediated signaling [38]. Arabidopsis’s serine/threonine phosphatase 7 (AtPP7) has been shown to act as a nuclear localization signaling molecule that dephosphorylates nuclear signaling intermediates (probably unknown) when exposed to blue light. It acts in the downstream of cryptochrome and in the response of blue light ensuring stable signaling [38].

#### 3.3.2 The output pathways regulated by core clock

Proteins Ote100001960091 and Ote100189530101 have been identified as Ethylene receptor 1 and ethylene response sensor 1 respectively. Both of which encode histidine kinases similar to members of two-component regulator family and have been shown to be integral components in ethylene signaling [8]. The gaseous hormone has a large number of implications like fruit-ripening, senescence, stress response, seed germination and many more and its signaling has been shown to be under circadian control [61]. Protein Ote100058660011 has been predicted as Gibberellin (GA) receptor GID1B that acts as a soluble gibberellin receptor. GA is a diterpenoid hormone that regulates various stages of plant growth and development like seed germination, development of fruit, pollen tube and flowering time. GID1 binds with biologically active GAs like GA1, GA3 and GA4 targets a GRAS family protein DELLA that negatively regulate GA signaling for proteasomal degradation [64]. Proteins Ote100237240051 and Ote100260840121 are predicted to be UDP-N-acetylglucosamine–peptide N-acetylglucosaminyltransferase SPINDLY (SPY) that negatively regulate Gibberellin signaling to inhibit hypocotyl elongation. It has been shown that SPY and GI function together and GI acts as a light gated negative regulator of SPY function to control various physiological processes like elongation of hypocotyl, flowering time and cotyledon movements [63]. Ote100003860081, predicted as Two-component response regulator ARR3, behaves like a bacterial two component system that negatively regulate cytokine signaling and affects the phosphorylation state of ARRs by transducing the signal through a multi-step His-Asp phosphotransfer process [62]. Ote100257670011 has been characterized as Histidine kinase 3 protein that acts as a positive regulator of cytokine signaling [40]. Proteins Ote100167620091 and Ote100120330051 have been predicted to be Two-component response regulator-like APRR1, APRR7 respectively. They are similar to ARR family His-Asp phosphorelay systems except that they the aspartate site that accept phosphate group is missing. Circadian control at transcriptional level of all the members of APRR1/TOC1 family has been documented and they have shown to express sequentially in order of APRR9, APRR7, APRR5, APRR3, and APRR1 within a period of 24*h* [30]. At dawn CCA1 and LHY negatively regulate APRR1, CHE and their own expression. But during the day CHE levels rise that repress CCA1 and finally at evening TOC1 reset the clock cycle by repressing the expression of CHE [45]. Similarly APRR7 has also been shown to repress CCA1 and LHY expressions and regulate expression of other clock components [39]. It acts as master regulator of various physiological processes of plant including response to abiotic stress [27], photoperiodic flowering and temperature compensation [39]. The results also confirmed the identities of predicted clock proteins and show that most of the proteins possess their respective functions (Table S2). Furthermore, protein family and domain databases integrated by InterProScan also confirmed our results (Table S4).

## 4 Summary and Conclusions

Considering the importance of circadian rhythms in regulating physiological processes of the plant, current study pursues the identification and characterization of genes involved in generating these oscillations in *O. tenuiflorum*. It will increase our understanding of various components of circadian circuitry responsible for various circadian rhythms as well as the underlying mechanisms sustaining these rhythms which is largely unknown in Tulsi. In this study we propose a methodology to identify 24 known core circadian clock genes in Tulsi that leverage in house built HMMs from 56 template plants. Further we propose a hybrid methodology that combines RWR and GDV local topological measures to identify novel proteins associated with core clock. Based on RWR, GDV and two statistical significance tests a total of 70 putative CC associated genes proteins were obtained. Functional annotation of the candidate proteins was performed and we found that a large number of proteins are involved in regulation of gene expression. We obtained crucial proteins transducing red, far-red and blue light signals to the core oscillator. Furthermore, we obtained proteins involved in different physiological process, like, gibberllin signaling, ethylene signlaing, cytokine signaling, abiotic and biotic stress response etc. We hope that the proposed framework for reporting genome-wide identification and characterization of core circadian clock and associated proteins in *O. tenuiflorum* shall pave the way for prioritizing the CC or any other biological process associated proteins in plants.

## Acknowledgments

VS^†^ thanks Council of Scientific and Industrial Research (CSIR), India for providing Junior Research Fellowship (JRF).

## Authors Contributions

VS^∗^ conceptualized and designed the research framework. VS^†^ performed the computational experiments. VS^†^ and VS^∗^ analyzed the data and interpreted results. VS^†^ and VS^∗^ wrote and finalized the manuscript.

## Data and materials availability

All data is available in the manuscript or the supplementary materials.

## References

[1] Genome sequencing of herb Tulsi (Ocimum tenuiflorum) unravels key genes behind its strong medicinal properties. BMC Plant Biology, 2015.

[2] David Alabadí, Tokitaka Oyama, Marcelo J Yanovsky, Franklin G Harmon, Paloma Más, and Steve A Kay. Reciprocal regulation between toc1 and lhy/cca1 within the arabidopsis circadian clock. Science, 293(5531):880–883, 2001.

[3] Stephen F Altschul, Warren Gish, Webb Miller, Eugene W Myers, and David J Lipman. Basic local alignment search tool. Journal of molecular biology, 215(3):403–410, 1990.

[4] Rolf Apweiler, Amos Bairoch, Cathy H Wu, Winona C Barker, Brigitte Boeckmann, Serenella Ferro, Elisabeth Gasteiger, Hongzhan Huang, Rodrigo Lopez, Michele Magrane, et al. Uniprot: the universal protein knowledgebase. Nucleic acids research, 32(suppl 1):D115–D119, 2004.

[5] Tanya Z Berardini, Leonore Reiser, Donghui Li, Yarik Mezheritsky, Robert Muller, Emily Strait, and Eva Huala. The arabidopsis information resource: making and mining the “gold standard” annotated reference plant genome. genesis, 53(8):474–485, 2015.

[6] Pedro Carmona-Saez, Monica Chagoyen, Francisco Tirado, Jose M Carazo, and Alberto Pascual-Montano. Genecodis: a web-based tool for finding significant concurrent annotations in gene lists. Genome biology, 8(1):1–8, 2007.

[7] Lei Chen, Chen Chu, Jing Lu, Xiangyin Kong, Tao Huang, and Yu-Dong Cai. A computational method for the identification of new candidate carcinogenic and non-carcinogenic chemicals. Molecular BioSystems, 11(9):2541–2550, 2015.

[8] Karen L Clark, Paul B Larsen, Xiaoxia Wang, and Caren Chang. Association of the arabidopsis ctr1 raf-like kinase with the etr1 and ers ethylene receptors. Proceedings of the National Academy of Sciences, 95(9):5401–5406, 1998.

[9] MarcMaurice Cohen. Tulsi - Ocimum sanctum: A herb for all reasons. Journal of Ayurveda and Integrative Medicine, 2014.

[10] Michael F Covington, Julin N Maloof, Marty Straume, Steve A Kay, and Stacey L Harmer. Global transcriptome analysis reveals circadian regulation of key pathways in plant growth and development. Genome biology, 9(8):1–18, 2008.

[11] Antony N Dodd, Neeraj Salathia, Anthony Hall, Eva Kévei, Réka Tóth, Ferenc Nagy, Julian M Hibberd, Andrew J Millar, and Alex AR Webb. Plant circadian clocks increase photosynthesis, growth, survival, and competitive advantage. Science, 309(5734):630–633, 2005.

[12] Zhou Du, Xin Zhou, Yi Ling, Zhenhai Zhang, and Zhen Su. agrigo: a go analysis toolkit for the agricultural community. Nucleic acids research, 38(suppl 2):W64–W70, 2010.

[13] Rachel S Edgar, Edward W Green, Yuwei Zhao, Gerben Van Ooijen, Maria Olmedo, Ximing Qin, Yao Xu, Min Pan, Utham K Valekunja, Kevin A Feeney, et al. Peroxiredoxins are conserved markers of circadian rhythms. Nature, 485(7399):459–464, 2012.

[14] Zheng Eelderink-Chen, Jasper Bosman, Francesca Sartor, Antony N Dodd, Ákos T Kovács, and Martha Merrow. A circadian clock in a nonphotosynthetic prokaryote. Science advances, 7(2):eabe2086, 2021.

[15] Kathleen Greenham and C Robertson McClung. Integrating circadian dynamics with physiological processes in plants. Nature Reviews Genetics, 16(10):598–610, 2015.

[16] Linqu Han, Mary Mason, Eddy P Risseeuw, William L Crosby, and David E Somers. Formation of an scfztl complex is required for proper regulation of circadian timing. The Plant Journal, 40(2):291–301, 2004.

[17] Stacey L Harmer, John B Hogenesch, Marty Straume, Hur-Song Chang, Bin Han, Tong Zhu, Xun Wang, Joel A Kreps, and Steve A Kay. Orchestrated transcription of key pathways in arabidopsis by the circadian clock. Science, 290(5499):2110–2113, 2000.

[18] Carlos T Hotta, Michael J Gardner, Katharine E Hubbard, Seong Jin Baek, Neil Dalchau, Dontamala Suhita, Antony N Dodd, and Alex AR Webb. Modulation of environmental responses of plants by circadian clocks. Plant, cell & environment, 30(3):333–349, 2007.

[19] Keisuke Inoue, Takashi Araki, and Motomu Endo. Integration of input signals into the gene network in the plant circadian clock. Plant and Cell Physiology, 58(6):977–982, 2017.

[20] Jose A Jarillo, Juan Capel, Ru-Hang Tang, Hong-Quan Yang, Jose M Alonso, Joseph R Ecker, and Anthony R Cashmore. An arabidopsis circadian clock component interacts with both cry1 and phyb. Nature, 410(6827):487–490, 2001.

[21] Philip Jones, David Binns, Hsin-Yu Chang, Matthew Fraser, Weizhong Li, Craig McAnulla, Hamish McWilliam, John Maslen, Alex Mitchell, Gift Nuka, et al. Interproscan 5: genome-scale protein function classification. Bioinformatics, 30(9):1236–1240, 2014.

[22] Minoru Kanehisa and Susumu Goto. Kegg: kyoto encyclopedia of genes and genomes. Nucleic acids research, 28(1):27–30, 2000.

[23] Woe-Yeon Kim, Sumire Fujiwara, Sung-Suk Suh, Jeongsik Kim, Yumi Kim, Linqu Han, Karine David, Joanna Putterill, Hong Gil Nam, and David E Somers. Zeitlupe is a circadian photoreceptor stabilized by gigantea in blue light. Nature, 449(7160):356–360, 2007.

[24] Sebastian Köhler, Sebastian Bauer, Denise Horn, and Peter N Robinson. Walking the interactome for prioritization of candidate disease genes. The American Journal of Human Genetics, 82(4):949–958, 2008.

[25] Philippe Lamesch, Tanya Z Berardini, Donghui Li, David Swarbreck, Christopher Wilks, Rajkumar Sasidharan, Robert Muller, Kate Dreher, Debbie L Alexander, Margarita Garcia-Hernandez, et al. The arabidopsis information resource (tair): improved gene annotation and new tools. Nucleic acids research, 40(D1):D1202–D1210, 2012.

[26] Gang Li, Hamad Siddiqui, Yibo Teng, Rongcheng Lin, Xiang-yuan Wan, Jigang Li, On-Sun Lau, Xinhao Ouyang, Mingqiu Dai, Jianmin Wan, et al. Coordinated transcriptional regulation underlying the circadian clock in arabidopsis. Nature cell biology, 13(5):616–622, 2011.

[27] Tiffany Liu, Jenny Carlsson, Tomomi Takeuchi, Linsey Newton, and Eva M Farre. Direct regulation of abiotic responses by the a rabidopsis circadian clock component prr 7. The plant journal, 76(1):101–114, 2013.

[28] Hua Lu, C Robertson McClung, and Chong Zhang. Tick tock: circadian regulation of plant innate immunity. Annual review of phytopathology, 55:287–311, 2017.

[29] Linyuan Lü, Duanbing Chen, Xiao-Long Ren, Qian-Ming Zhang, Yi-Cheng Zhang, and Tao Zhou. Vital nodes identification in complex networks. Physics Reports, 650:1–63, 2016.

[30] Akinori Matsushika, Seiya Makino, Masaya Kojima, and Takeshi Mizuno. Circadian waves of expression of the aprr1/toc1 family of pseudo-response regulators in arabidopsis thaliana: insight into the plant circadian clock. Plant and Cell Physiology, 41(9):1002–1012, 2000.

[31] C Robertson McClung. Plant circadian rhythms. The Plant Cell, 18(4):792–803, 2006.

[32] C Robertson McClung. Beyond arabidopsis: the circadian clock in non-model plant species. 24(5):430–436, 2013.

[33] Todd P Michael and C Robertson McClung. Enhancer trapping reveals widespread circadian clock transcriptional control in arabidopsis. Plant physiology, 132(2):629–639, 2003.

[34] Todd P Michael, Todd C Mockler, Ghislain Breton, Connor McEntee, Amanda Byer, Jonathan D Trout, Samuel P Hazen, Rongkun Shen, Henry D Priest, Christopher M Sullivan, et al. Network discovery pipeline elucidates conserved time-of-day–specific cis-regulatory modules. PLoS genetics, 4(2):e14, 2008.

[35] Tijana Milenković and Nataša Pržulj. Uncovering biological network function via graphlet degree signatures. Cancer informatics, 6:CIN–S680, 2008.

[36] Joel C Miller and Aric Hagberg. Efficient generation of networks with given expected degrees. In International Workshop on Algorithms and Models for the Web-Graph, pages 115–126. Springer, 2011.

[37] Lalit Mohan, M. V. Amberkar, and Meena Kumari. Ocimum sanctum linn (TULSI) - an overview, 2011.

[38] Simon G Møller, Youn-Sung Kim, Tim Kunkel, and Nam-Hai Chua. Pp7 is a positive regulator of blue light signaling in arabidopsis. The Plant Cell, 15(5):1111–1119, 2003.

[39] Norihito Nakamichi, Takatoshi Kiba, Rossana Henriques, Takeshi Mizuno, Nam-Hai Chua, and Hitoshi Sakakibara. Pseudo-response regulators 9, 7, and 5 are transcriptional repressors in the arabidopsis circadian clock. The Plant Cell, 22(3):594–605, 2010.

[40] Chika Nishimura, Yoshi Ohashi, Shusei Sato, Tomohiko Kato, Satoshi Tabata, and Chiharu Ueguchi. Histidine kinase homologs that act as cytokinin receptors possess overlapping functions in the regulation of shoot and root growth in arabidopsis. The Plant Cell, 16(6):1365–1377, 2004.

[41] Maria A Nohales and Steve A Kay. Molecular mechanisms at the core of the plant circadian oscillator. Nature structural & molecular biology, 23(12):1061–1069, 2016.

[42] Abhay Kumar Pandey, Pooja Singh, and Nijendra Nath Tripathi. Chemistry and bioactivities of essential oils of some Ocimum species: an overview. Asian Pacific Journal of Tropical Biomedicine, 4(9):682–694, sep 2014.

[43] Prachi Pandey, Vadivelmurugan Irulappan, Muthukumar V Bagavathiannan, and Muthappa Senthil-Kumar. Impact of combined abiotic and biotic stresses on plant growth and avenues for crop improvement by exploiting physio-morphological traits. Frontiers in plant science, 8:537, 2017.

[44] Priyabrata Pattanayak, Pritishova Behera, Debajyoti Das, and SangramK Panda. Ocimum sanctum Linn. A reservoir plant for therapeutic applications: An overview. Pharmacognosy Reviews, 2010.

[45] Jose L Pruneda-Paz, Ghislain Breton, Alessia Para, and Steve A Kay. A functional genomics approach reveals che as a component of the arabidopsis circadian clock. Science, 323(5920):1481–1485, 2009.

[46] Nataša Pržulj. Biological network comparison using graphlet degree distribution. Bioinformatics, 23(2):e177–e183, 2007.

[47] Natasa Pržulj, Derek G Corneil, and Igor Jurisica. Modeling interactome: scale-free or geometric? Bioinformatics, 20(18):3508–3515, 2004.

[48] Shubhra Rastogi, Alok Kalra, Vikrant Gupta, Feroz Khan, Raj Kishori Lal, Anil Kumar Tripathi, Sriram Parameswaran, Chellappa Gopalakrishnan, Gopalakrishna Ramaswamy, and Ajit Kumar Shasany. Unravelling the genome of Holy basil: An “incomparable” “elixir of life” of traditional Indian medicine. BMC Genomics, 2015.

[49] Shubhra Rastogi, Seema Meena, Ankita Bhattacharya, Sumit Ghosh, Rakesh K. Shukla, Neelam S. Sangwan, Raj K. Lal, Madan M. Gupta, Umesh C. Lavania, Vikrant Gupta, Dinesh A. Nagegowda, and Ajit K. Shasany. De novo sequencing and comparative analysis of holy and sweet basil transcriptomes. BMC Genomics, 2014.

[50] Shubhra Rastogi, Saumya Shah, Ritesh Kumar, Divya Vashisth, Md Qussen Akhtar, Ajay Kumar, Upendra Nath Dwivedi, and Ajit Kumar Shasany. Ocimum metabolomics in response to abiotic stresses: Cold, flood, drought and salinity. PloS one, 14(2):e0210903, 2019.

[51] Jong Sang Ryu, Jeong-Il Kim, Tim Kunkel, Byung Chul Kim, Dae Shik Cho, Sung Hyun Hong, Seong-Hee Kim, Aurora Piñas Fernández, Yumi Kim, Jose M Alonso, et al. Phytochrome-specific type 5 phosphatase controls light signal flux by enhancing phytochrome stability and affinity for a signal transducer. Cell, 120(3):395–406, 2005.

[52] David J Sheerin, Chiara Menon, Sven zur Oven-Krockhaus, Beatrix Enderle, Ling Zhu, Philipp Johnen, Frank Schleifenbaum, York-Dieter Stierhof, Enamul Huq, and Andreas Hiltbrunner. Light-activated phytochrome a and b interact with members of the spa family to promote photomorphogenesis in arabidopsis by reorganizing the cop1/spa complex. The Plant Cell, 27(1):189–201, 2015.

[53] Fabian Sievers, Andreas Wilm, David Dineen, Toby J Gibson, Kevin Karplus, Weizhong Li, Rodrigo Lopez, Hamish McWilliam, Michael Remmert, Johannes Söding, et al. Fast, scalable generation of high-quality protein multiple sequence alignments using clustal omega. Molecular systems biology, 7(1):539, 2011.

[54] E Singh and S Sharma. Diversified potentials of Ocimum sanctum Linn (Tulsi): An exhaustive survey. J Nat Prod Plant Resour, 2(1):39–48, 2012.

[55] Gagandeep Singh, Vikram Singh, and Vikram Singh. Genome-wide interologous interactome map (teagpin) of camellia sinensis. Genomics, 113(1):553–564, 2021.

[56] Gagandeep Singh, Vikram Singh, and Vikram Singh. Systems scale characterization of circadian rhythm pathway in camellia sinensis. Computational and Structural Biotechnology Journal, 2022.

[57] RajKumar Brojen Singh, Vikram Singh, and Ram Ramaswamy. Stochastic synchronization of circadian rhythms. Journal of Systems Science and Complexity, 23(5):978–988, 2010.

[58] Vikram Singh, Gagandeep Singh, and Vikram Singh. Tulsipin: an interologous protein interactome of ocimum tenuiflorum. Journal of proteome research, 19(2):884–899, 2019.

[59] David E Somers, Paul F Devlin, and Steve A Kay. Phytochromes and cryptochromes in the entrainment of the arabidopsis circadian clock. Science, 282(5393):1488–1490, 1998.

[60] Deepti Srivastava, Md Shamim, Mahesh Kumar, Anurag Mishra, Rashmi Maurya, Divakar Sharma, Pramila Pandey, and KN Singh. Role of circadian rhythm in plant system: An update from development to stress response. Environmental and Experimental Botany, 162:256–271, 2019.

[61] Simon C Thain, Filip Vandenbussche, Lucas JJ Laarhoven, Mandy J Dowson-Day, Zhi-Yong Wang, Elaine M Tobin, Frans JM Harren, Andrew J Millar, and Dominique Van Der Straeten. Circadian rhythms of ethylene emission in arabidopsis. Plant Physiology, 136(3):3751–3761, 2004.

[62] Jennifer PC To, Georg Haberer, Fernando J Ferreira, Jean Deruere, Michael G Mason, G Eric Schaller, Jose M Alonso, Joseph R Ecker, and Joseph J Kieber. Type-a arabidopsis response regulators are partially redundant negative regulators of cytokinin signaling. The Plant Cell, 16(3):658–671, 2004.

[63] Tong-Seung Tseng, Patrice A Salomeé, C Robertson McClung, and Neil E Olszewski. Spindly and gigantea interact and act in Arabidopsis thaliana pathways involved in light responses, flowering, and rhythms in cotyledon movements. The Plant Cell, 16(6):1550–1563, 2004.

[64] Miyako Ueguchi-Tanaka, Motoyuki Ashikari, Masatoshi Nakajima, Hironori Itoh, Etsuko Katoh, Masatomo Kobayashi, Teh-yuan Chow, Yueie C Hsing, Hidemi Kitano, Isomaro Yamaguchi, et al. Gibberellin insensitive dwarf1 encodes a soluble receptor for gibberellin. Nature, 437(7059):693–698, 2005.

[65] Wei Wang, Jinyoung Yang Barnaby, Yasuomi Tada, Hairi Li, Mahmut Tör, Daniela Caldelari, Dae-un Lee, Xiang-Dong Fu, and Xinnian Dong. Timing of plant immune responses by a central circadian regulator. Nature, 470(7332):110–114, 2011.

[66] Esther Yakir, Dror Hilman, Yael Harir, and Rachel M Green. Regulation of output from the plant circadian clock. The FEBS journal, 274(2):335–345, 2007.

[67] Ö mer Nebil Yaveroğlu, Noël Malod-Dognin, Darren Davis, Zoran Levna- jic, Vuk Janjic, Rasa Karapandza, Aleksandar Stojmirovic, and Nataša Pržulj. Revealing the hidden language of complex networks. Scientific reports, 4(1):1–9, 2014.

[68] Jia Ye, Lin Fang, Hongkun Zheng, Yong Zhang, Jie Chen, Zengjin Zhang, Jing Wang, Shengting Li, Ruiqiang Li, Lars Bolund, et al. Wego: a web tool for plotting go annotations. Nucleic acids research, 34(suppl 2):W293–W297, 2006.

[69] Shai Yerushalmi and Rachel M Green. Evidence for the adaptive significance of circadian rhythms. Ecology letters, 12(9):970–981, 2009.

[70] Jae-Woong Yu, Vicente Rubio, Na-Yeoun Lee, Sulan Bai, Sun-Young Lee, Sang-Sook Kim, Lijing Liu, Yiyue Zhang, María Luisa Irigoyen, James A Sullivan, et al. Cop1 and elf3 control circadian function and photoperiodic flowering by regulating gi stability. Molecular cell, 32(5):617–630, 2008.

